# Unilateral damage to the entopeduncular nucleus causes forelimb motor dysfunction in rats

**DOI:** 10.1101/2025.05.30.657131

**Authors:** Ryo Sakai, Kazuki Kuroda, Takashi Ryoke, Ayako Maegawa, Koshi Murata, Yugo Fukazawa

## Abstract

**Background:** The entopeduncular nucleus (EP), corresponding to the human globus pallidus internal segment, is a basal ganglia output nucleus, and plays a critical role in motor control. However, the impact of EP damage on skilled motor function and the relationship between its damage in stroke, such as internal capsule hemorrhage (ICH), and motor dysfunction remains unclear. This study aimed to clarify whether EP damage causes motor dysfunction in two disease models.

**Methods:** EP-related motor dysfunction was investigated by inducing localized unilateral EP damage in Long-Evans rats using a stereotactic kainic acid (KA) injection. Motor function was assessed using a single-pellet reaching task pre-injection and on postoperative days 2, 7, 14, 21, and 28. Immunohistochemical staining for NeuN, somatostatin (SST), and parvalbumin was conducted to quantify damage and its correlation with motor outcomes. In addition, unilateral ICH was induced via stereotactic injection of collagenase type IV, which dissolves the vascular basement membrane, into the internal capsule (IC) of Long-Evans rats. Injury sites were classified into the IC, dorsomedial region from the IC, ventral lateral region from the IC, and EP, and their volumes were measured. Measured volumes were analyzed for correlations with motor function assessments.

**Results:** KA-induced EP damage significantly reduced reaching success rates on postoperative day 2 compared to those in the control group (p<0.05). Immunohistochemical analysis showed that reaching success rates on day 28 positively correlated with the numbers of remaining NeuN-positive and SST-positive neurons (p<0.05). In the ICH experiment, all rats significantly reduced the success rate of the reaching task to 0% on day 2, and the success rate on day 28 correlated positively with the remaining EP volume, but not with total lesion volume.

**Conclusions:** EP damage was strongly associated with motor impairments, highlighting its critical role in motor control and recovery.

## Introduction

The entopeduncular nucleus (EP), equivalent to the human globus pallidus internal segment, is an output nucleus of the basal ganglia, playing a critical role in motor control.^1,2^ Striatal dopamine receptor D1 and D2 neurons directly and indirectly, respectively, project to EP neurons in mice, and optogenetic manipulation of individual striatal neurons causes complex disruptions in learned forelimb movements, suggesting a role of EP in the integration of motor-related signals from striatum.^3^ In rodent models of Parkinson’s disease, nigral dopamine depletion alters the activity of neurons in EP as well as that of striatum and subthalamic nucleus, which results in motor dysfunction.^4^ Studies using animal models of Parkinson’s disease have demonstrated that optogenetic inhibition of EP neuronal activity improves ipsilateral rotational movements and stepping behavior.^5,6^ These findings strongly suggest that EP plays a pivotal role in motor control and its damage could be involved in neurological diseases accompanying motor dysfunction.

The physiological role of EP in motor function and the pathophysiological impact of EP damage in neurological diseases remain elusive. Previous studies on motor dysfunction with subcortical hemorrhage in rats have demonstrated that lesions in the posterior ventral internal capsule (IC) result in the most severe motor impairments.^7–10^ Notably, the posterior ventral part of the IC includes the EP, suggesting that EP damage exacerbates motor dysfunction severity in these studies. Several studies have investigated the direct effects of EP damage on motor function, but their results remain controversial. Studies inducing EP lesions with ibotenic acid have shown no significant impacts on rotarod tests and locomotor activity.^11–13^ In contrast, that with muscimol induced bradykinesia and hypometria during reaching movements.^14^ Recent studies have revealed that EP consists of several subtypes of neurons in the mouse EP.^15,16^ At least four distinct neuronal subtypes were identified in EP, with parvalbumin (PV)-positive neurons and somatostatin (SST)-positive neurons accounting for more than 70% of the total population of EP neurons.^17^ These two neuronal subtypes exhibit distinct distributions within EP, with PV-positive neurons predominantly located in the caudal region and SST-positive neurons in the rostral region.^17^ As prior studies have focused on global lesion volume for the lesion assessment without evaluating the loss of EP neuron subtypes,^11,13,18^ the inconsistency in motor dysfunction caused by EP lesions could be attributed to the variability in the damage (loss) of neuronal types within EP. Thus, conducting a detailed quantitative assessment of the loss of individual neuron types would provide further insights into the physiological role of EP on motor function and the pathophysiological impact of EP damage in neurological diseases accompanying motor dysfunction, including stroke.

Rat models of stroke are widely used to study the mechanisms underlying motor dysfunction after stroke.^19,20^ Among these, the collagenase-induced intracerebral hemorrhage (ICH) model is superior to the traditional blood injection model, better mimicking the pathological condition caused by vascular wall damage.^19,21^ Additionally, behavioral tests that allow detailed assessment of pathological motor states after the stroke onset are essential to analyze changes in motor function, and recent studies using rat stroke models have employed the single-pellet reaching task.^22–25^ The reaching movement of rats in this task is structurally similar to human reaching movements, and it has been reported to reproduce human stroke-related motor dysfunction severity.^26,27^ Thus, combining the ICH model with reaching movement analysis would provide an ideal condition to elucidate the impact of EP damage on motor function and the mechanisms underlying functional impairments by stroke.

In this study, we investigate whether selective EP lesions and ICH affect motor function by assessing the reaching movement before and after the operations. This study will provide a comprehensive knowledge of the pathophysiology underlying motor dysfunction after stroke, and establish a robust framework for future investigations into the mechanisms driving motor impairments associated with EP damage.

## Methods

### Animals

Experiments were conducted under the Guidelines for Animal Experimentation in Neuroscience proposed by the Japan Neuroscience Society and approved by the Experimental Animal Research Committee of the University of Fukui (R05014) and Fukui Health Science University (R01-001). Male Long-Evans rats (Sankyo Labo Service Corporation, Inc., Japan) were housed under a 12-h light/dark cycle, with food and water available *ad libitum,* including during the training period for the single-pellet reaching task. Intracranial microinjections were performed in 24 rats (Figures 1–3), of which 5 received saline and 19 received kainic acid (KA), including 6 rats used for optimizing injection coordinates. Of the KA-injected rats, 5 were excluded from the analysis due to no tissue damage identified via Nissl staining (Figure S1). Nissl staining confirmed that the damage in the included KA-injected rats was confined to the EP judged by aggregates of degenerating cells, with no detectable neuronal damage in the putamen or thalamus. For ICH experiments, 24 rats were used to optimize the concentration and dose of collagenase and the coordinates of the intracranial microinjection, and another 11 rats were used to obtain the behavioral and histological data depicted in Figures 4–6.

**Figure 1.**
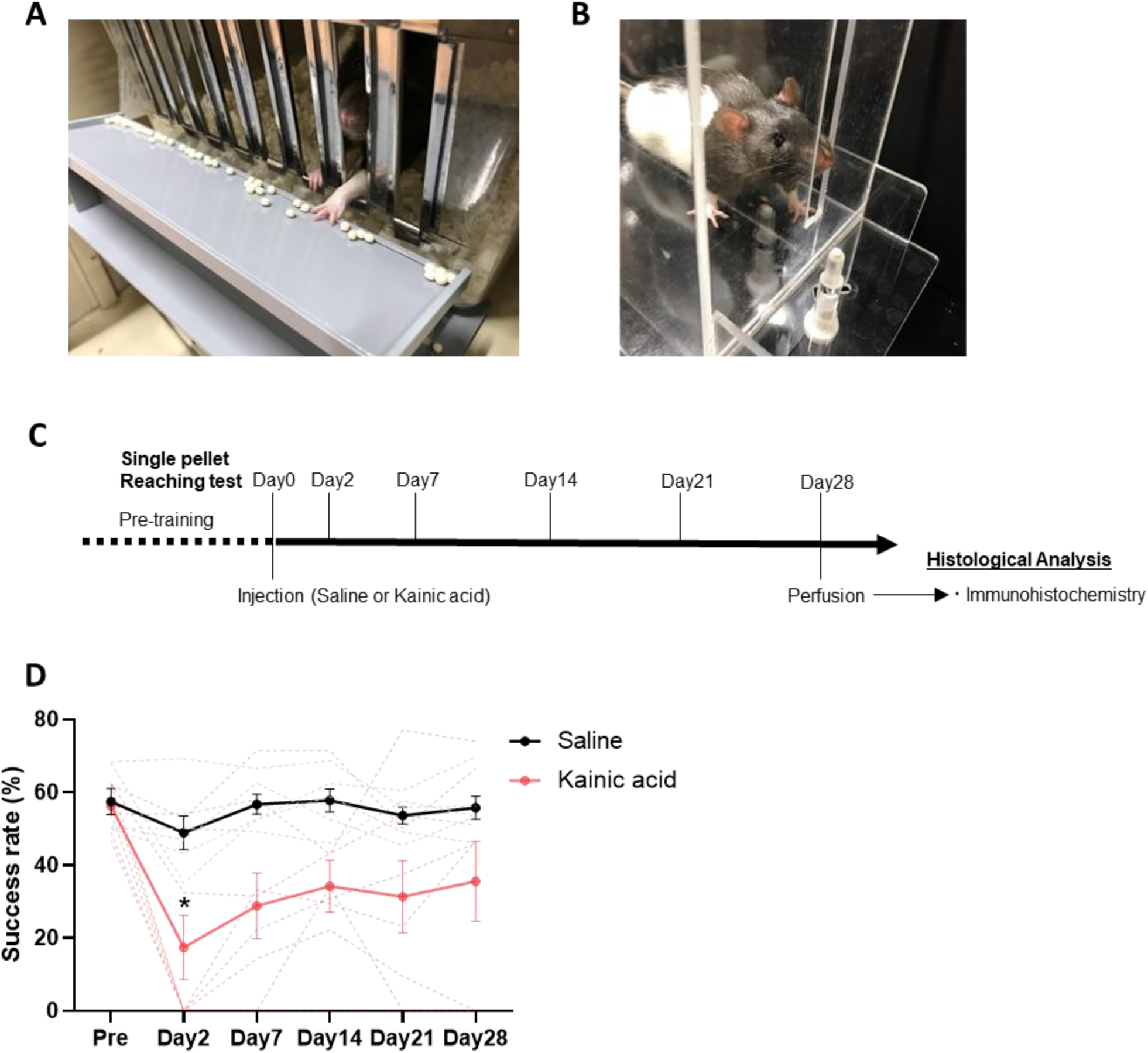
Evaluation of motor function after unilateral injection of KA in the EP. A. Home cage equipped with food trays. The food tray is positioned equivalent to the reaching distance used in the single-pellet reaching task. The dominant limb of each rat is identified after approximately 1 week of training using the food tray. B. Single-pellet reaching box. The feeding table on which the pellet is placed can be moved in a V-shape. Therefore, conditions can be changed according to the dominant limb, and only the dominant limb can be used. C. Experimental schedule of behavioral assessments using the single-pellet reaching task after a single injection of KA. The task was performed preoperatively and on days 2, 7, 14, 21, and 28 post-injection. D. Results of the single-pellet reaching task. Solid lines indicate the mean, and dotted lines represent individual results. Data are presented as the mean ± SEM (n=5 for saline, n=8 for KA). Statistical significance was calculated using a two-way mixed-effects model, followed by Bonferroni’s multiple comparisons test. *p<0.05. EP, entopeduncular nucleus; SEM, standard error of mean; ANOVA, analysis of variance

### EP damage model by KA administration

Rats underwent operation at 10–12 weeks of age (350–450 g body weight [B.W.]). The animals were anesthetized with a mixture of medetomidine (0.4 mg/kg B.W.), midazolam (2.0 mg/kg B.W.), and butorphanol (5.0 mg/kg B.W.). The head was fixed to a stereotaxic frame (SR-6R-HT; Narishige Co., Ltd., Japan), and the scalp was cut along the midline. A small hole was drilled 3.8 mm lateral to the midline and 2.0 mm caudal to the bregma, contralateral to the dominant forelimb determined during the single-pellet reaching task training. A total of 0.3 μL of KA (diluted to 2.5 µg/µL with sterile saline; Sigma-Aldrich, USA) or saline alone was injected at a rate of 0.1 µL/min over 7 min using a glass micropipette that was connected to a Hamilton syringe at a depth of 8.0 mm below the dura mater. The needle was held in place for 7 min after the injection and then slowly pulled upwards. The holes in the skull were covered with circular plastic disks and fixed using dental acrylic resin. The wound was sutured, and antisedan (1.2 mg/kg B.W. atipamezole) was administered intraperitoneally. Rats were kept warm on a heat pad until they awakened.

### ICH model

The surgical protocol for the ICH model with forelimb dysfunction was based on previous studies with optimized injection coordinates for Long-Evans rats.^28,29,24^ As in the KA administration experiment, rats were operated on at 10–12 weeks of age (350–450 g B.W.). A small hole was drilled at 3.8 mm lateral to the midline and 2.0 mm caudal to the bregma, contralateral to the preferred forelimb. To induce local hemorrhage at the IC, a total of 1.4 µL collagenase type Ⅳ (15 U/mL in sterile saline; Sigma-Aldrich), which digests the vascular basement membrane, was injected at a rate of 0.2 µL/min over 7 min at a depth of 5.8 mm below the dura mater. The needle was held in place for 7 min after the injection and then slowly pulled out.

### Behavioral assessments

The single-pellet reaching task was conducted as described previously.^3,22,30–33^ Before the preoperative behavioral test, rats were trained to reach food pellets (45 mg, Bioserv) for approximately 1 week in their home cages (height, 25 cm; width, 33 cm; length, 38 cm). The home cage had 1.3 cm wide slits, with a food tray placed 1.8 cm away, allowing the rats to reach the pellets by extending their forelimbs (Figure 1A). Rats had free access to standard chow in addition to food pellets (2 g/day) replenished once daily.

After the home cage training, reaching behavior was assessed in a single-pellet reaching box (height, 47.5 cm; width, 13 cm; length, 45 cm) (Figure 1B). An adjustable food pole (height from the chamber floor, 4 cm; diameter, 7 mm) was placed 0.5–1.5 cm away from a slit (height, 15 cm; width, 1.3 cm), and pellets were placed one at a time in an indentation at the top of the pole. Rats were placed in the box and trained to reach a pellet through the slit and to retrieve a pellet placed on the pole diagonally to the preferred forelimb. The reaching test was conducted daily for 20 min or until the rats caught 30 pellets. Performance was scored according to the success rate, as follows: success rate (%) = (number of times the rat caught pellets/number of times the forelimb was extended toward the pellet from the slit) × 100. Rats were subjected to the microinjection surgery when the success rate of the single-pellet reaching task was >50%. The single-pellet reaching task was performed preoperatively and at 2, 7, 14, 21, and 28 days postoperatively.

### Histological assessment of damage induced by KA administration

After the behavioral test on postoperative day 28, the rats were deeply anesthetized via intraperitoneal injection of sodium pentobarbital (100 mg/kg B.W, Nacalai Tesque, Japan), perfused with phosphate-buffered saline (PBS, 0.14 M NaCl, 0.027 M KCl, 0.033 M Na₂HPO₄, and 0.002 M KH₂PO₄; pH 7.4),, and perfusion-fixed with 4% paraformaldehyde in 0.1M phosphate buffer. Rat brains were carefully removed and immersed in the same fixative solution overnight for post-fixation.

For histological assessment of damage by Nissl staining and immunohistochemistry (IHC), the fixed brains were subjected to sucrose substitution, freeze-embedded in OCT compound (Tissue-Tech), and sliced into 20-µm sections using a cryostat. A series of sections were collected at 200-µm intervals and stained by Nissl staining for neuronal loss, or immunolabelled with specific antibodies to analyze the neuronal subtypes remaining after KA administration.

Sections stained by Nissl staining were examined using a bright-field virtual slide system (NanoZoomer; Hamamatsu Photonics, Shizuoka, Japan). Images were digitized with NanoZoomer, cropped using ImageJ (National Institutes of Health), and reconstructed in 3D using the “Reconstruct” software^34^ to calculate the area and extent of damage on each slide.

In IHC, sections were labeled with anti-substance P antibody to define EP region and with anti-NeuN antibody to identify neurons surviving after KA administration. Remaining neurons were classified into subtypes based on SST and PV expression. Immunostaining was performed by rinsing sections three times in PBS and briefly immersed in PBS. Antigen retrieval was conducted using HistoVT (diluted 1:10; Nacalai Tesque, Kyoto, Japan) at 70°C for 20 min. The sections were rinsed three times in PBS, permeabilized in TNT (0.1 M Tris-HCl, pH 7.5; 0.15 M NaCl; 0.1% Tween 20) for 15 min, and blocked with 10% normal donkey serum (NDS) diluted in TNT for 30 min at room temperature. The sections were then incubated overnight at 4 °C with the following primary antibodies diluted in 0.2% NDS-TNT: mouse anti-NeuN (1:500, Sigma-Aldrich), goat anti-PV (1:500, Nittobo Medical Co., Ltd, Japan), rabbit anti-SST (1:500, GeneTeX, USA), and rat anti-substance P (1:500, Sigma-Aldrich) antibodies. On the second day, the sections were washed twice in TNT for 5– 10 min each and incubated for 1 h at room temperature with fluorescent dye-conjugated secondary antibodies diluted in 0.2% NDS-TNT: Cy3-conjugated anti-mouse (1:500), Alexa Fluor 488-conjugated anti-goat (1:500, Jackson ImmunoResearch, USA), Cy3-conjugated anti-rabbit (1:500, Jackson ImmunoResearch), and Alexa Fluor 647-conjugated anti-rat (1:500, Jackson ImmunoResearch) antibodies. After two washes in TNT for 5 min each, the sections were counterstained with 4’,6-diamidino-2-phenylindole (1:1000 dilution in PBS) for 5 min at room temperature. Finally, the sections were washed once in PBS for 10 min and mounted using ProLong Glass (Thermo Fisher Scientific, USA). IHC images were captured using an all-in-one fluorescence microscope (BZ-X800, KEYENCE, Osaka, Japan). The EP area on each slide and the number of cells within that area were quantified using Fiji (an ImageJ distribution, National Institutes of Health, USA). The EP area was defined based on SP labeling. This comprehensive approach enabled the analysis of both overall tissue damage and detailed neuronal subtype classification.

### Histological assessment of damage and its location in ICH model rats

The brains of ICH model rats were collected by the same procedure as in the KA administration experiment explained above. To determine the area of hemorrhage in the ICH model rats, brains were sectioned coronally at 100-µm thickness using a vibratome, and a series of sections at 100-µm intervals were collected on glass slides and air-dried overnight. The sections were delipidated in ethanol, and stained with hematoxylin and eosin (HE).^35^ After dehydration, the sections were cleared with xylene and sealed with a coverslip using mounting medium, and then examined as in the KA experiment.

The damaged area was classified based on the outline of the IC, with defined dorsal-medial and ventral-lateral outflow. The remaining EP area was manually tracked. The residual EP volume was estimated as follows: the contralateral EP was traced, and the percentage of remaining EP on the injection side was calculated by dividing the EP volume on the damaged side by the EP volume on the contralateral side. The slice containing the maximal anterior commissure width was set as 1.0 mm from the bregma, and the coordinates of the damaged area were calculated.^36^

### Data analysis

Data for the success rate in the single-pellet reaching task, the damaged area by KA or ICH and cell counts were collected by different experimenters who were unaware of the experimental groups. All statistical analyses were performed using GraphPad Prism 8 (GraphPad Software Inc., USA). Normality was assessed using the Shapiro–Wilk test prior to all analyses. To evaluate the effects of group (saline vs. KA) and time on reaching performance in the KA model experiment, a two-way mixed-effects model was applied, followed by Bonferroni’s multiple comparisons test. Depending on data distribution, either the Mann–Whitney U test or unpaired t-test was used to compare results of Nissl staining and IHC between the saline and KA groups. Histological analyses (IHC and HE staining) and the results of the reaching task were assessed using Pearson’s correlation coefficient or Spearman’s rank correlation coefficient, depending on data distribution. Significant correlations were further evaluated using simple linear regression. For the ICH model experiment, one-way analysis of variance followed by Tukey’s post hoc test was used to compare time-point data from the single-pellet reaching task. Data are presented as mean ± standard error of the mean, and statistical significance was set at p < 0.05.

## Results

### Localized KA-induced damage to the EP impaired single-pellet reaching performance

Motor deficits of the forelimbs were evaluated using the single-pellet reaching task following unilateral KA injection into the EP. The reaching task was performed preoperatively (3–6 h before surgery, [day 0]) and on days 2, 7, 14, 21, and 28 postoperatively (Figure 1C). Rats in the KA group showed a significantly lower success rate than those in the saline group on postoperative day 2 (Figure 1D and Video S1), whereas no significant difference in success rate was observed between the saline and KA groups on postoperative days 7, 14, 21 and 28. These results suggest that the unilateral injection of KA into the EP rapidly leads to transient motor dysfunction.

To examine the loss of EP neurons induced by KA injection, immunostaining for NeuN was performed, and SP was co-stained to define the EP region (Figure 2A).^15^ The area of EP showed no significant difference between the saline and KA groups (p=0.1063, Figure 2B). The number of NeuN-positive cells within the EP region, however, was significantly reduced in the KA group compared to that in the saline group (p <0.01, Figure 2C). A decrease in the number of NeuN-positive cells was observed throughout the EP, ranging from bregma -2.8 to -3.2 mm (Figure 2D), confirming widespread neuronal loss in the EP by KA injection.

**Figure 2.**
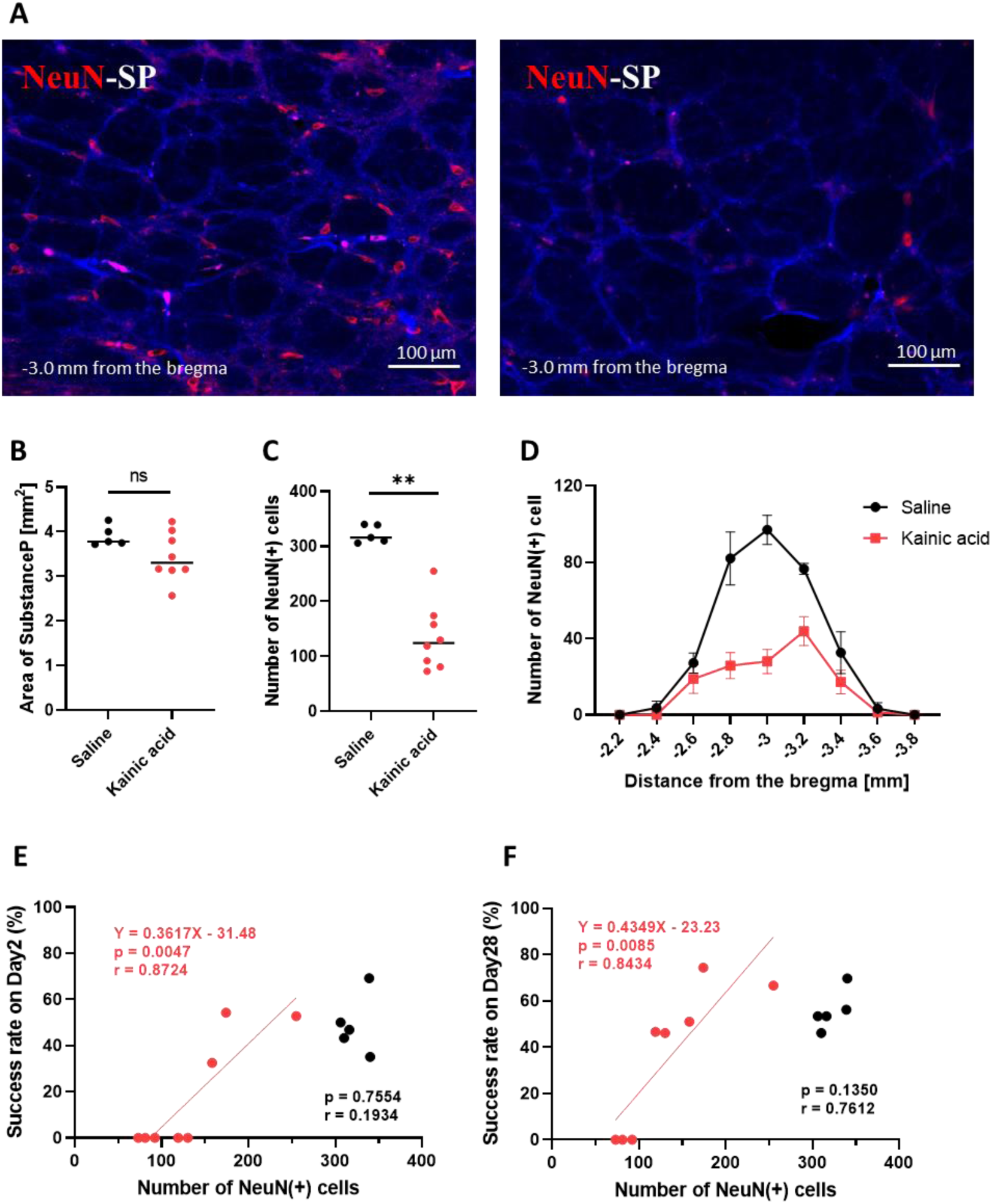
Quantitative analysis of neuronal damage in the EP. A. Immunohistochemical staining results of NeuN (red) and SP (blue). B. Comparison of SP+ regions. C. Comparison of NeuN+ cell counts. D. Distribution of damaged NeuN+ cells along the rostrocaudal axis. E, F. Relationship between the number of NeuN+ cells and performance in the single-pellet reaching task (E, day 2; F, day 28). Data are presented as the mean ± SEM (n=5 for saline, n=8 for KA). Statistical significance was determined using unpaired t-test and Pearson’s correlation coefficient. **p<0.01. EP, entopeduncular nucleus; SP, substance P; SEM, standard error of mean

The correlation between the loss of EP neurons and motor dysfunction and recovery following EP damage was investigated. On postoperative days 2 and 28, the number of NeuN-positive cells positively correlated with the success rate (day 2; r=0.8724, p<0.01, day 28; r=0.8434, p<0.01, Figure 2E, F). Notably, among the eight rats, five and three rats with the lowest number of residual NeuN-positive cells showed 0% success rate on postoperative day 2 and 28, respectively. These results suggest that localized damage to the EP impairs single-pellet reaching performance and that severe EP damage may lead to persistent motor dysfunction.

EP neurons comprise several cell types.^15–17,37^ To gain insights into which EP neuron cell types were damaged by KA injection and whether motor dysfunction correlates with a loss of specific neuron types, the numbers of residual SST-positive and PV-positive neurons, two major cell types in the EP, were evaluated (Figure 3A). IHC revealed a significant reduction in the number of both neuron types in the KA group compared to that in the saline group (p<0.01, Figure 3B), with more robust loss of SST-positive neurons than PV-positive neurons. To investigate whether these subtypes influence motor dysfunction following EP damage, the correlation between the success rate on days 2 and 28 and the number of each cell type was analyzed. A significant positive correlation was observed between the number of SST-positive neurons and the success rate on both day 2 and day 28 (day 2, r = 0.7958, p < 0.05; day 28, r = 0.8468, p < 0.05), whereas no significant correlation was found for PV-positive neurons (day 2, r = 0.4637, p = 0.2619; day 28, r = 0.7075, p = 0.0640) (Figure 3C). To examine the anatomical characteristics of these cell types in more detail, we further analyzed the rostro-caudal distribution of SST-positive and PV-positive neurons in the EP, as well as their reduction by KA injection along the rostrocaudal axis. We confirmed that SST-positive neurons were predominantly distributed in the rostral region of the EP, whereas PV-positive neurons were primarily distributed in the caudal region, consistent with a previous report on mice (Figure 3D, E).^15–17,37^ The KA injection reduced the numbers of both SST-positive and PV-positive neurons over the rostrocaudal axis.

**Figure 3.**
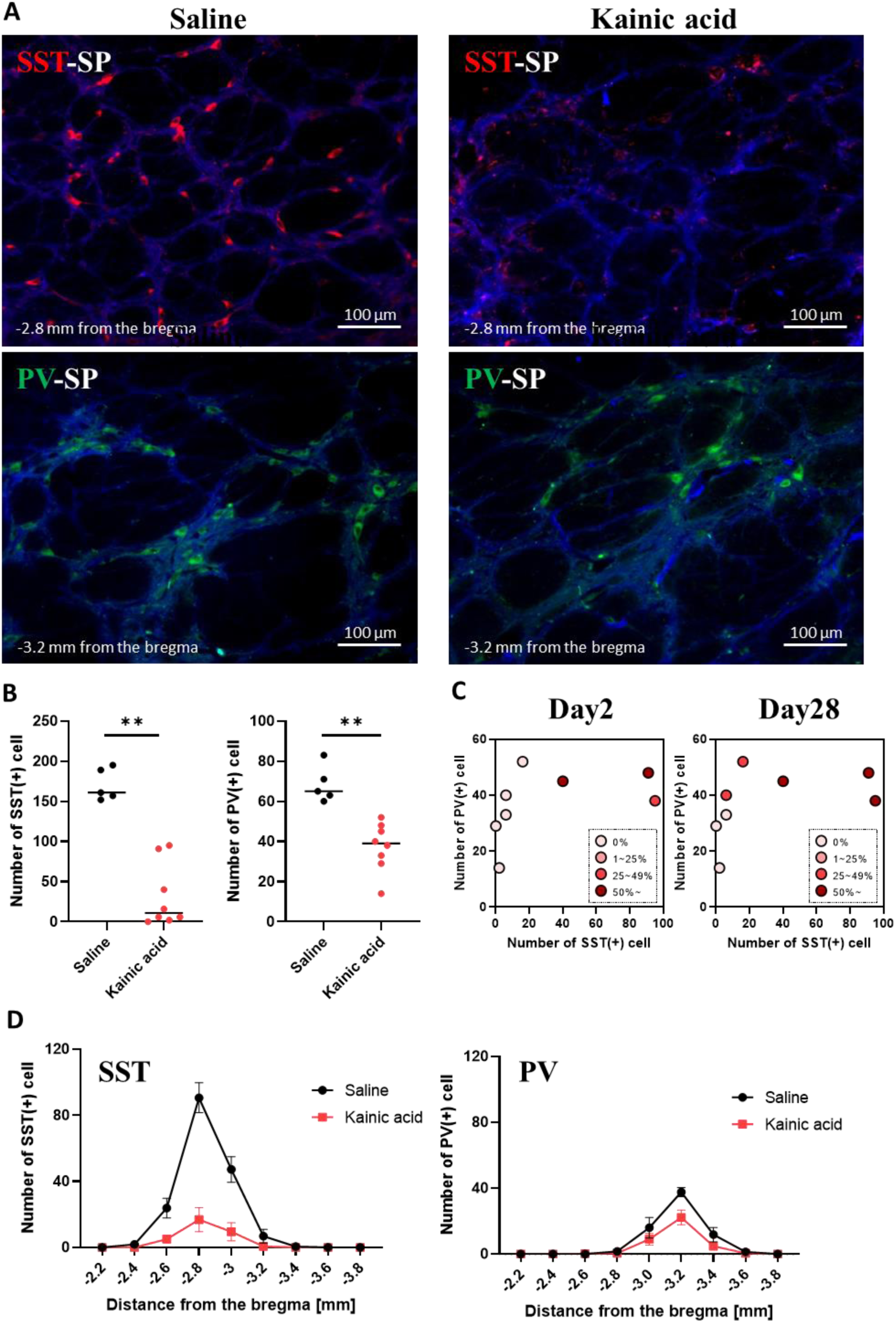
Quantitative analysis of SST+ neurons in the EP following KA administration. A. Immunohistochemical staining results of SST (red), PV (green), and SP (blue). B. Comparison of cell counts of SST+ and PV+ cells. C. Visualization of the relationship between the SST+ and PV+ cell counts and motor impairments in individual subjects on days 2 and 28. D. Distribution of damaged SST+ and PV+ cells along the rostrocaudal axis. Data are presented as the mean ± SEM (n=5 for saline, n=8 for KA). Statistical significance was determined using the Mann–Whitney U test (SST) and unpaired t-test (PV). **p<0.01. EP, entopeduncular nucleus; SP, substance P; SST, somatostatin; PV, parvalbumin; SEM, standard error of mean

Taken together, our data suggest that unilateral KA injection into the EP causes death of both SST-positive and PV-positive neurons and induces acute and persistent motor dysfunction.

### Collagenase-induced ICH impaired single-pellet reaching performance

To verify the influence of EP damage in a subcortical hemorrhage, hemorrhage was induced by injecting collagenase into the IC region near the EP. The schedule for the single-pellet reaching task and injection surgery was similar to that for the KA administration experiments: preoperative day 0 and postoperative days 2, 7, 14, 21, and 28 (Figure 4A). The preoperative trial success rate was 60.3±2.9 (n=11 rats). On postoperative day 2, no rats succeeded in the reaching task (0.0%). The success rate was significantly lower at all postoperative time points compared to the preoperative day (day 0 vs. day 2, p<0.0001; day 0 vs. day 7, p=0.0001; day 0 vs. day 14, p=0.0001; day 0 vs. day 21, p=0.0002; day 0 vs. day 28, p=0.0026, Figure 4B). Although the average success rate on day 28 remained lower than the preoperative success rate, four of the 11 rats had a 0% success rate, whereas the 7 showed varying success rates, ranging from 5% to 60%.

**Figure 4.**
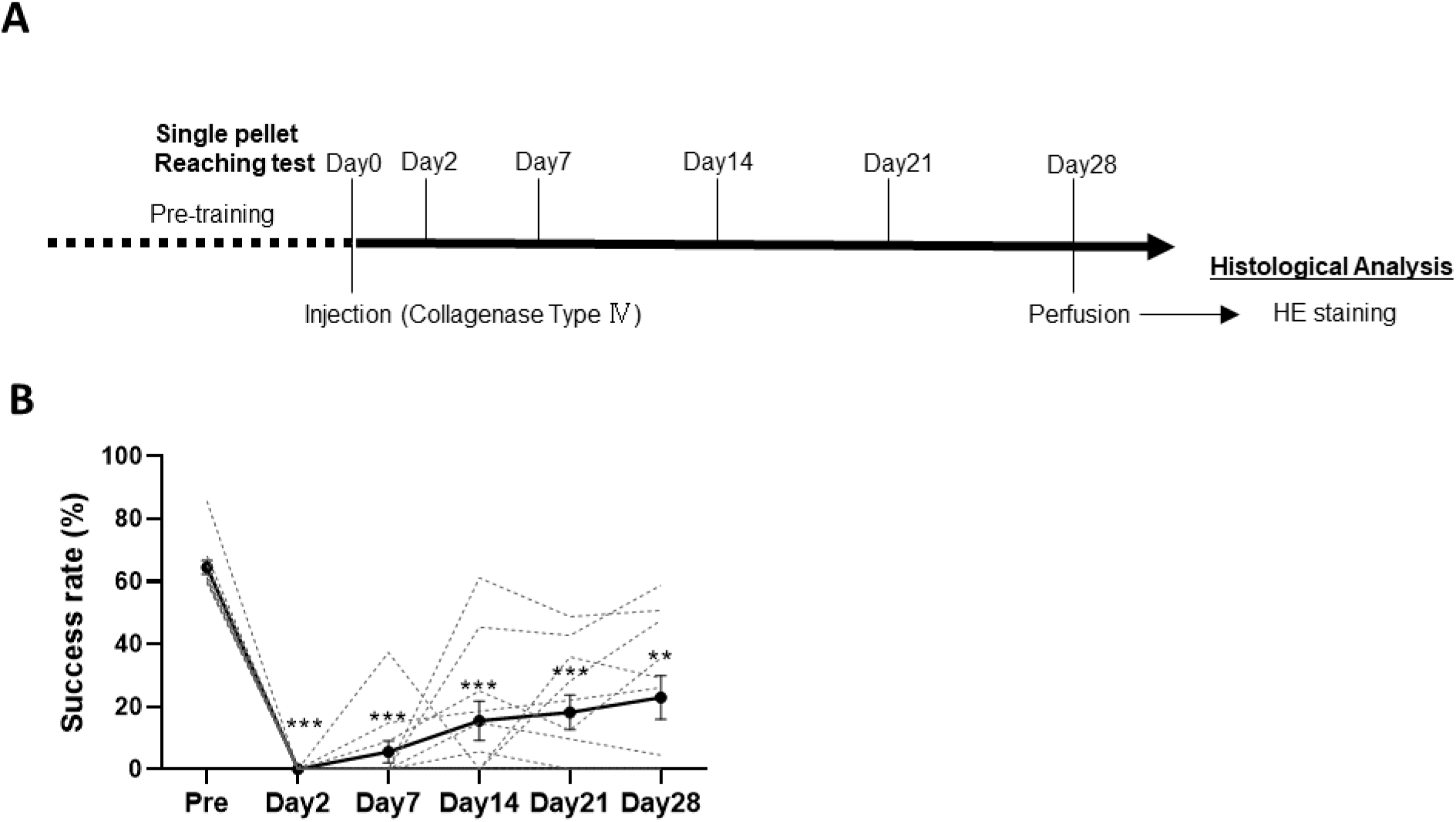
Evaluation of motor function in ICH model rats. A. Experimental schedule of behavioral assessments using the single-pellet reaching task after a single unilateral collagenase injection. The task was performed preoperatively and on days 2, 7, 14, 21, and 28 after the injection. B. Results of the single-pellet reaching task. Black solid lines indicate the mean, and gray dotted lines represent individual results. Data are presented as the mean ± SEM (n=11). Statistical significance was calculated using a one-way ANOVA with post-hoc Tukey’s test. **p<0.01, ***p<0.001. ICH, internal capsule hemorrhage; HE, hematoxylin and eosin; SEM, standard error of mean; ANOVA, analysis of variance

### Motor recovery after ICH correlated with Residual EP volume

Histological analysis revealed that collagenase injection successfully induced ICH in all 11 rats. The IC is a white-matter structure composed of bundles of myelinated fibers, partially surrounding the EP. Notably, the extent of hemorrhage in the EP largely varied among individual rats (Figure 5A). Therefore, we investigated whether motor recovery on day 28 (Figure 4B) correlated with EP hemorrhage volume. As hemorrhage obscured the histological landmarks of the EP, making direct assessment of the lesional area difficult, the percentage of the residual EP volume was used as an index (68.5±8.05%, Figure 5A). The success rate of the single-pellet reaching task on day 28 positively correlated with the residual EP volume (r=0.7997, p<0.01, Figure 5B). Notably, the residual EP volume was not dependent on the total lesion volume (p=0.4758, Figure 5C). These data suggest that the extent of EP lesions is a pivotal factor in motor recovery after ICH.

**Figure 5.**
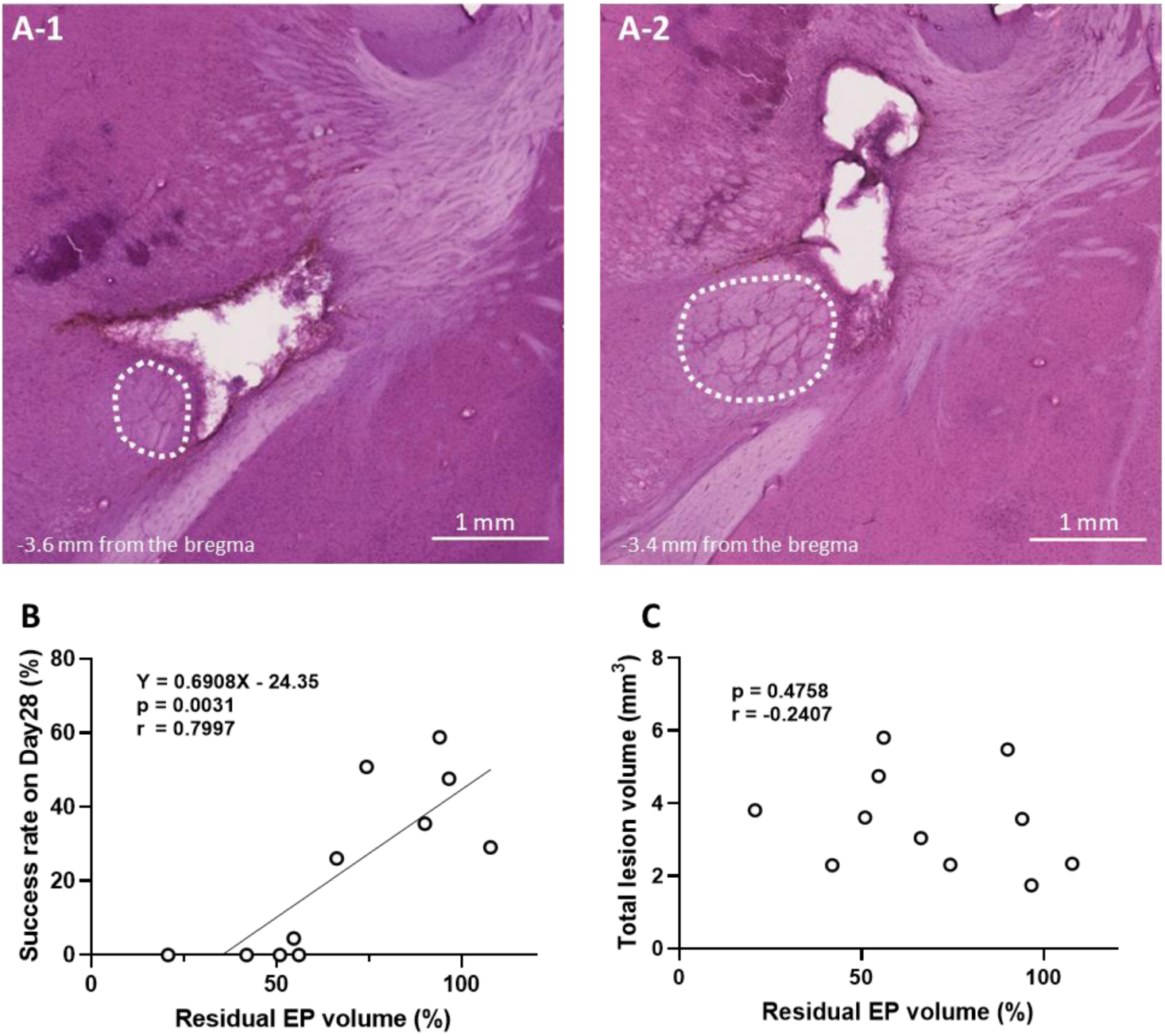
Relationship between residual EP volume and behavior tests on postoperative day 28. A. Examples of the damaged area of the EP. A-1. Most of the EP is damaged. A-2. Most of the EP is not damaged. B. Relationship between the results of the single-pellet reaching task test on postoperative day 28 and residual EP volume. C. Relationship between residual EP volume and total lesion volume. Statistical significance was determined using the Pearson’s correlation coefficient. Scale bars: 1 mm. EP, entopeduncular nucleus

### Lesioned volumes in IC and surrounding regions do not correlate with motor deficits

To evaluate the possibility that the motor dysfunction in ICH was caused by the damage in the surrounding regions of IC, damage to the IC and surrounding regions was measured. Although hemorrhage primarily occurred in the IC, some rats also exhibited hemorrhage and tissue damage in regions outside the IC. The injured regions were classified as follows: the IC, ventrolateral (VL) region extending to the striatum, and dorsomedial (DM) region extending to the thalamus (Figure 6A, B). The average injury volumes in the regions were as follows: IC, 2.7±0.26; VL, 0.6±0.25; DM, 0.2 ± 0.07; and total lesion volume, 3.5±0.40 mm^3^ (Figure 6C, D). We then investigated whether recovery of reaching test scores on day 28, the time point showing large individual differences (Figure 6E), correlated with the volume of the hemorrhage area for each region. No statistically significant correlation was observed between the lesion volume in any region (IC, VL, DM, or entire lesion) and behavioral scores on day 28 (success rate of the single-pellet reaching task vs. IC, r=-0.2777, p=0.4084; vs. VL, r=-0.3236, p=0.3316; vs. DM r=0.03776, p=0.9122; vs. total, r=-0.3710, p=0.2625, Figure 6E).

**Figure 6.**
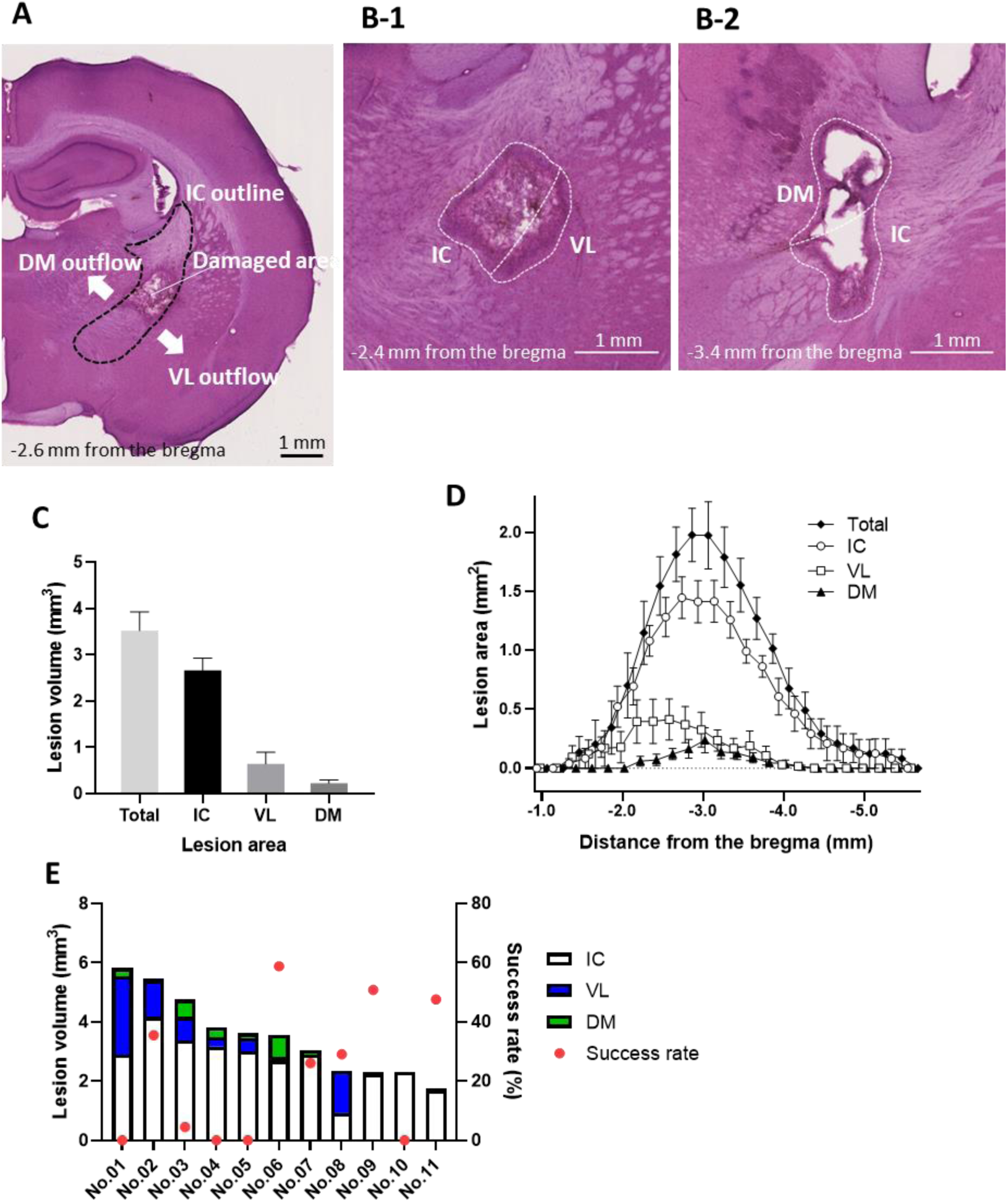
Characterization of damaged brain areas in ICH rats on day 28 after the collagenase injection. A. Representative hematoxylin-eosin (HE) stained images of brain slices obtained from ICH rats. The black dotted line outlines the internal capsule (IC). White arrows indicate the damaged area that extends beyond the IC. Dorsomedial (DM) and ventrolateral (VL) outflow regions are defined. B. Typical examples of damage classification. B-1: IC and VL, B-2: IC and DM. C. Total volume of the entire lesion, IC, VL, and DM. D. Distribution of the damaged area along the rostrocaudal axis. E. Relationship between the results of the single-pellet reaching task on day 28 and total volume of the entire lesion, IC, VL, or DM. Data are presented as the mean ± SEM (n=11). ICH, internal capsule hemorrhage; SEM, standard error of mean

## Discussion

The impact of EP damage on motor function remains elusive. A previous study reported that bilateral EP lesions induced via intracranial microinjection of ibotenate did not affect locomotor function in rats.^11^ In contrast, in a human case report of unilateral damage to the globus pallidus, the importance of its internal segment was highlighted in the pathophysiology of dystonia.^38^ Another study administering muscimol to the internal segment of the globus pallidus in macaque monkeys reported bradykinesia and low-amplitude forelimb movements.^14^ In addition, lesions in the IC and EP are reported to result in long-lasting impairments of skilled motor function in a rat model of photothrombotic infarct.^9^ Collectively, the damage of EP is likely to lead to motor impairments that are distinct from those caused by corticospinal tract damage. However, these studies lacked anatomical characterization of the damage to the EP and surrounding regions, and thus, the effect of EP damage alone on motor function has remained elusive. In the present study, we quantitatively assessed damage to the EP and surrounding regions in two lesion models in rats and correlated it with dysfunction in the single pellet reaching task to explore how the EP damage affects skilled motor function and its recovery.

In the EP damage model, KA was locally injected into the EP unilaterally, which induces excitotoxicity in neurons via binding to ionotropic glutamate receptors expressed on the somato-dendritic region of neurons. This approach selectively damaged local EP neurons while sparing surrounding areas, and resulted in acute forelimb motor dysfunction in most of KA-injected animals. Moreover, a positive correlation between the number of remaining EP neurons and reaching test scores on day 28 was observed, substantiating a pivotal role of EP in motor function. To further confirm whether EP damage causes motor dysfunction in a subcortical hemorrhage model, the histological changes of the EP and surrounding areas were compared with the severity of forelimb motor dysfunction in the rat ICH model. Hemorrhage at the IC caused severe and long-lasting motor dysfunction, as previously demonstrated.^39,40,41^ Due to the uncontrollable nature of the hemorrhage, large inter-individual variability was observed in the volume and location of damage and the motor dysfunction outcomes. None of the rats with IC hemorrhage succeeded in the single-pellet reaching task on postoperative day 2, but the scores on day 28 varied among individuals. No significant correlation was observed between the damage volume of the IC or surrounding areas (such as striatum and thalamus) and reaching test scores. On the other hand, the volume of EP damage significantly correlated with reaching test scores on day 28, similar to the results observed in the KA model. These findings demonstrate that the EP plays a pivotal role in skilled motor function, and that its damage causes motor dysfunction, the recovery of which is also dependent on the remaining amount of the EP itself.

EP neuron cell types are less defined in rats compared to in mice, in which 3 types of projection neurons, glutamate/GABA co-releasing SST-positive neurons, glutamatergic PV-positive neurons, and GABAergic PV-positive neurons, have been classified and PV-positive and SST-positive neurons account for more than 70% of the total population in the EP.^42–44^ Our examination revealed the existence of these two types of neurons in the rat EP and their rostrocaudal distribution pattern similar to those of mice: SST-positive neurons were predominantly distributed in the rostral region, whereas PV neurons were concentrated in the caudal region. In KA-injected rats, the two neuron types were reduced to different extents, leading to variability in motor dysfunction following EP lesions. These results support the hypothesis that distinct neuron types within the EP contribute uniquely to motor function, and that the observed variability in motor deficits reflects the differential loss of these specific neuron types. Notably, a significant positive correlation was observed between the number of remaining SST-positive neurons and reaching performance. However, studies by Rajakumar et al.^45^ and Wallace et al.^44^ have reported that PV-positive neurons, unlike SST-positive neurons that project mainly to the lateral habenula, send denser projections to motor-related thalamic areas such as the ventro-anterior lateral, ventro-medial, anterodorsal thalamus, parafascicular nucleus, and brainstem. In line with the anatomical findings, the present study revealed that rats with substantial SST-positive neuron loss exhibited only mild motor impairment, whereas others with less PV-positive neuron loss demonstrated marked dysfunction (Figure 3C). This finding suggests that the loss of PV-positive neurons may have a greater impact on motor function. Additionally, the EP has been shown to project to the accessory olivary nucleus, suggesting functional connections with cerebellar and other motor-related regions via circuits outside the classical basal ganglia loop.^46^ Given that KA-induced lesion employed in this study affects both SST-positive and PV-positive neurons regardless of the projection targets, further investigation by cell-type– and projection-specific neuronal manipulations is imperative to determine which EP neuron types are critically implicated in skilled motor function.

Compensatory neuroplasticity involving the ipsilateral and contralateral corticospinal tracts, as well as the corticorubral and corticoreticular pathways, is known to play a critical role in motor recovery following pyramidal tract injury.^47^ In contrast, the mechanisms underlying motor recovery following the EP damage remain largely unknown. In the present study, a significant positive correlation was observed between the extent of EP damage and the degree of functional recovery (Figure 2F), suggesting the existence of compensatory mechanisms for motor restoration after EP damage and the importance of the EP in the restoration. Further investigation of the recovery mechanisms following EP damage would provide novel insights into the neuroplasticity mechanisms underlying functional recovery after a subcortical hemorrhage.

## Acknowledgments

We thank Eri Murai, Takako Maegawa, Mayumi Yamamoto, Meiko Hagihara, Yuichiro Ishiyama, Mari Onoi, Risa Takada and members of the Fukazawa Laboratory at the University of Fukui and Tokuichi Iguchi, Toui Takagi, Azusa Ideguchi, and Jun Ishimaru at Fukui Health Science University for their technical assistance. We also thank Hideki Yoshikawa for fabricating the devices used for behavioral analysis.

## Supplemental Material

Figure S1

Video S1

## Non-standard Abbreviations and Acronyms

EP: entopeduncular nucleus
GABA: gamma-aminobutyric acid
IC: internal capsule
ICH: internal capsule hemorrhage
IHC: immunohistochemical staining
KA: kainic acid
PBS: phosphate-buffered saline
PV: parvalbumin
SP: substance P
SST: somatostatin

